# “Remote adipose tissue-derived stromal cells of patients with lung adenocarcinoma generate a similar malignant microenvironment of the lung stromal counterpart”

**DOI:** 10.1101/2022.07.08.499306

**Authors:** Elena De Falco, Antonella Bordin, Cecilia Menna, Xhulio Dhori, Paolo Rosa, Erino Rendina, Mohsen Ibrahim, Antonella Calogero

## Abstract

Cancer alters both local and distant tissue by influencing the microenvironment. In this regard, the interplay with the stromal fraction is considered critical as this latter can either foster or hamper the progression of the disease. Accordingly, the modality by which tumor may alter distant niches of stromal cells is still unclear, expecially at early stages. In this short report we attempt to better understand the biology of this cross talk. In our “autologous stromal experimental setting”, we found that remote adipose tissue-derived mesenchymal stem cells (mediastinal AMSC) obtained from patients with lung adenocarcinoma sustain proliferation and clonogenic ability of A549 and human primary lung adenocarcinoma cells similarly to the autologous stromal lung counterpart (LMSC). This effect is not observed in lung benign diseases as the hamartochondroma. This finding was validated by conditioning benign AMSC with supernatants from LAC up to 21 days. The new reconditioned media of the stromal fraction so obtained, was able to increase cell proliferation of A549 cells at 14 and 21 days similarly to that derived from AMSC of patients with lung adenocarcinoma. The secretome generated by remote AMSC revealed overlapping to the corresponding malignant microenvironment of the autologous local LMSC. Among the plethora of 80 soluble factors analysed by arrays, a small pool of 5 upregulated molecules including IL1-β, IL-3, MCP-1, TNF-α and EGF, was commonly shared by both malignant-like autologous A- and LMSC derived microenvironments vs those benign. The bioinformatic analysis both revealed that these proteins were strictly and functionally interconnected to the lung fibrosis and proinflammation and that miR-126, 101, 486 and let-7-g were their main targets. Accordingly, we found that in lung cancer tissues and blood samples from the same set of patients here employed, miR-126 and miR-486 displayed the highest expression levels in tissue and blood, respectively.

## 1. Introduction

Mesenchymal stem cells (MSC) have been described as adult multipotent stem cells, showing many relevant properties, spanning from the ability to immunomodulate and migrate to specific sites of injury to the trans-differentiation into multiple cell types [1,2]. MSC have been considered ideal candidates for many clinical and cell therapy applications, almost concluding that their wide applicability was also possible in cancer treatment [3].

The biological interaction between MSC and tumor is complex and enormously debated. Several controversies exist about the potential of MSC to enhance or to even arrest tumorigenicity, because of their double faced behaviour: tumor-tropism (hence tested as vehicles for anticancer genes targeting cancer cells or as enhancement of the CAR-T immunotherapy)[4,5] and immunomodulative features, but also pro-metastatic functions [6-8], trans-differentiation into cancer-associated fibroblasts and drug resistance [9] and the parallel ability to overturn the immune system [10-13], activation of autophagy and neo-angiogenesis [14], therefore contributing to tumor evolution. This discrepancy also includes exosomes-derived MSC, considered both an intriguing therapeutic tool for drug delivery and main biological mediators of several supporting tumor molecular process [15]. Moreover, from a clinical standpoint, it has been recognized that the endogenous recruitment of MSC (of different origin including adipose) from systemic niches may occur by tumor secretion of inflammatory soluble factors [16] and that a correlation exists between circulating mesenchymal tumor cells and stage of tumor development [17,18].

This scenario is also complicated by recent indications about the heterogeneity of MSC and the phenotypic and functional changes potentially caused by tumors. For instance, adipose tissue and bone marrow derived MSC have shown differences with respect to stem cell content and epigenetic states [19,20]. Besides, MSC obtained from diverse sources such as heart, dermis, bone marrow and adipose tissue have been reported as genotypically different, expressing different levels of embryonic stem cell markers such as OCT-4, NANOG and SOX-2 [21], and biological properties including angiogenesis and secretome [20]. When MSC are derived from cancer tissues, they show altered molecular and functional properties [22-24], suggesting that the tumor characteristics such as benignity or malignancy could influence the environment where MSC are located.

From a biological standpoint, the evolution from a local to a systemic cancer microenvironment can be driven either by phenotypically altered cancer-associated cells (fibroblasts and endothelial cells), which organize clusters of systemic spreading cells or by niche-to niche recruiting phenomena from the bone marrow to the tumor site [25,26]. However, the thorny question is still centred on the modality by which cancer can control the systemic environment, influencing remote “normal” and non-bone marrow stem cell derived niches including distant MSC-niches, particularly at early stages of the tumor, which are of paramount biological and clinical relevance to understand cancer progression.

Assuming that the pathophysiology of cancer can be interpreted as a systemic disease, in this short report we attempt to investigate whether MSC-derived microenvironments at remote site from tumor can be already altered at early stages of lung adenocarcinoma.

## 2. Methods

### 2.1 Surgical specimen collection and clinical database

At the end of surgical procedure, a small sample of mediastinal adipose [1,2] and lung tissue was collected by electrocoagulation from patients undergoing surgical procedures for hamartochondroma and non-small cell lung carcinoma (NSCLC). Surgical procedures were conducted at S. Andrea Hospital, Rome. Written informed consent were obtained from patients, before starting all the surgical and laboratory procedures. Patients with NSCLC and staging T1N0M0 G1 were selected, whereas subjects with metastasis have been excluded from the study.

### 2.2 Isolation and Characterization of AMSC and LMSC

AMSC were isolated and characterized as previously described [27,1,2]. Lung specimens were chopped with scalpel and scissors in a 100mm Petri dish, then gently transferred into a clean 100mm Petri dish to allow tissue adherence. Complete growth medium composed by DMEM high glucose (Invitrogen) supplied by 10% FBS, antibiotics and L-glutamine (all Gibco) was added to the fragments. Plates were incubated at 37°C in a fully humidified atmosphere of 5% CO_2_, avoiding shaking the plates at least for 72 hours. Half of the medium was replaced with fresh complete medium every three days.

### 2.3 Isolation of lung adenocarcinoma cells and in vitro conditioning with MSC-derived supernatants

Human primary lung adenocarcinoma cells (LAC) were isolated as we already previously described [28]. The cells obtained were cultured in complete medium (DMEM-F12, penicillin-streptomycin, L-glutamine, nonessential aminoacids, sodium pyruvate, all Gibco, Monza, Italy and 5% FBS, Lonza, Milan, Italy). Lung adenocarcinoma A549 was purchased by ATCC and cultured in DMEM-F12 supplemented with 10% FBS (All Gibco). AMSC and LMSC supernatants derived from patients with hamartochondroma or NSCLC were collected between passage 3-6, then stored at -80°C until use. LAC cultures or A549 were conditioned by removing the medium and replacing with AMSC and LMSC supernatants at the following time course: 0, 3, 5, 7 days or 14 and 21 days for reconditioning of AMSC derived from patients with benign disease.

### 2.4 Proliferation and clonogenic assay

Both LAC and A549 were seeded onto 96 wells plates (150 cells each well) and incubated for 24 hours with DMEM low glucose 10% FBS [28]. Then cells were exposed to the different conditioned media collected from AMSC and LMSC up to 7 days. Cells treated with basal medium were used as control. The effect of conditioned media on cell viability was evaluated by the MTS assay. Briefly at 3, 5 and 7 days after treatment, 20μl of MTS reagent were added to each microculture well, and plates incubated for 2 hours at 37°C, after which absorbance at 492 nm (optical density) was measured using a microplate reader.

To test secondary colony forming efficiency (CFU) assay, LAC or A549 were seeded at passage 3 at low density (10 cells/cm^2^) [28] in AMSC and LMSC-derived conditioned media for 14 days and incubated at 37 °C. Colonies produced were fixed with 4% paraformaldehyde and then stained with Giemsa (Sigma, Milan, Italy) for 1 h and counted by optical microscope. A cluster with > 50 cells was considered as a colony [20].

### 2.5 Analysis of the autologous A- and LMSC-derived secretome

The evaluation of the different microenvironments was performed on collected supernatant obtained from both A- and LMSC in patients with lung adenocarcinoma and hemartochondroma. C-Series Human Cytokine Antibody Array C5 (RayBiotech, Inc) was used for simultaneous semi-quantitative detection of 80 multiple cytokines/growth factors as previously described [29]. Briefly, an equal volume of collected undiluted supernatants was incubated by gentle shaking overnight at 4°C on membrane of the kit C-Series Human Cytokine Antibody Array C5. Chemiluminescence was employed to quantify the spots (the same time exposure was used for all membranes) and each spot signal was analyzed by ImageJ. The samples were normalized on positive control means (six spots each array) and then values were expressed as percentage. To visualize the overall changes on cytokines array average data, results were graphed as log (2) in heatmap analysis by using the pheatmap R package in RdYlBl colour scale (from 0 to 7 expression levels). Cytokines with zero values in all replicates were ruled out.

### 2.6 Interaction and functional evaluation of cytokines network within the microenvironments and miRNA target interaction analysis

Correlation analysis between the cytokines expressed by A- or LMSC-derived benign and malignant microenvironments, was obtained by calculating the fold changes between these two conditions. A fold change threshold of >1.2 was considered as upregulated [30]. The analysis for known protein interaction was performed on common up-regulated cytokines between A- and LMSC by using the STRING software (https://string-db.org/, version 11.5) [31], building the whole network according to the high confidence setting (0.7) and default options. The pathway and process enrichment analysis were performed using Metascape as already described elsewhere [32]. The miRNA-target interaction analysis was performed by using “multiMiR” R package and review literature employing lung cancer as keyword. List of miRNAs in R package was obtained using get_multir function setting only on validated data and databases queried were miRecords, miRTarBase and TarBase.

### 2.7 MiRNA extraction and quantification

MicroRNAs extraction was performed from paraffin tumor tissue sections (RNeasy DSP FFPE Kit, Qiagen) and the RNA amount was determined using a Nanodrop spectrophotometer. Differently, miRNAs obtained from serum patients (200 µl) were isolated by the Qiagen miRNeasy kit with further modifications for biofluids applications. Syn-cel-miR-39 spike in synthetic RNA (Qiagen) was added to monitor extraction efficiency. Afterwards, on both tissue sections and sera samples the reverse transcription was performed using the “MiRCURY LNA Reverse Transcription Kit” (QIAGEN) in ThermoMixer 5436 (Eppendorf, Italy) according to the following protocol: 42°C for 60 seconds, 95°C for 5 minutes, 4°C forever [31].

Selected miRNA levels such as hsa-miR-101, hsa-miR-126, hsa-miR-486 and hsa-Let-7g were quantified by relative quantification using Qiagen LNA based SYBR green detection method (miRCURY LNA miRNA PCR assay-Qiagen). Briefly, 3μl of cDNA was used on the Applied Biosystems 7900HT machine, adding the relevant ROX concentration to the qPCR mix [31]. The relative miRNA expression was calculated using hsa-miR-16 as endogenous control with the 2-ΔCt method for cancer tissues and miR-16/miR-39 for sera [31].

### 2.8 Statistical Analysis

The results were expressed as the arithmetic mean ± standard deviation (SD) for at least 3 repeated individual experiments for each sample group. Statistical difference between the values were determined by student’s T-test, with a value of P < 0.05 was considered statistically significant.

## 3. Results

Firstly, we investigated if early-stage lung adenocarcinoma could influence the biological behaviour of MSC according to: 1) their own tissue source (autologous lung or mediastinal adipose tissue-derived MSC isolated from a tumor-free nearby or remote area, respectively); and 2) the biological feature of the lung tumour (malignant or benign defined by the histological analysis) where the MSC were derived from. To this aim, we conditioned the A549 cell line with supernatants of autologous lung or adipose-derived MSC (LMSC or AMSC) derived in parallel from patients with benign disease (pulmonary hamartochondroma) or malignant tumour (early-stage lung adenocarcinoma). We tested both cell proliferation (at 0, 3, 5, 7 days) and clonogenic capacity of A549. The experimental plan is depicted in Figure 1A. Results showed a significant increase of A549 cell proliferation at 5 and 7 days compared to control (Figure 1A, p<0.001 both vs control) after conditioning the lung tumor cell line with supernatants derived from LMSC or AMSC. When A549 were cultured with supernatants derived from autologous LMSC or AMSC both obtained from patients with pulmonary hamartochondroma, A549 proliferation decreased compared to control (Figure 1C, day 7 p<0.05 both *vs* control). Interestingly, a similar scenario was also reproduced regarding the clonogenic capacity of A549, where only mediastinal AMSC derived from patients with early-stage lung adenocarcinoma were able to enhance the clonogenicity respect to control (Figure 1C, p<0.05 *vs* control). The conditioning of A549 with supernatants derived from L- or AMSC of patients with pulmonary hamartochondroma, did not alter the clonogenic capacity of A549 (Figure 1C).

**Figure 1A-D.**
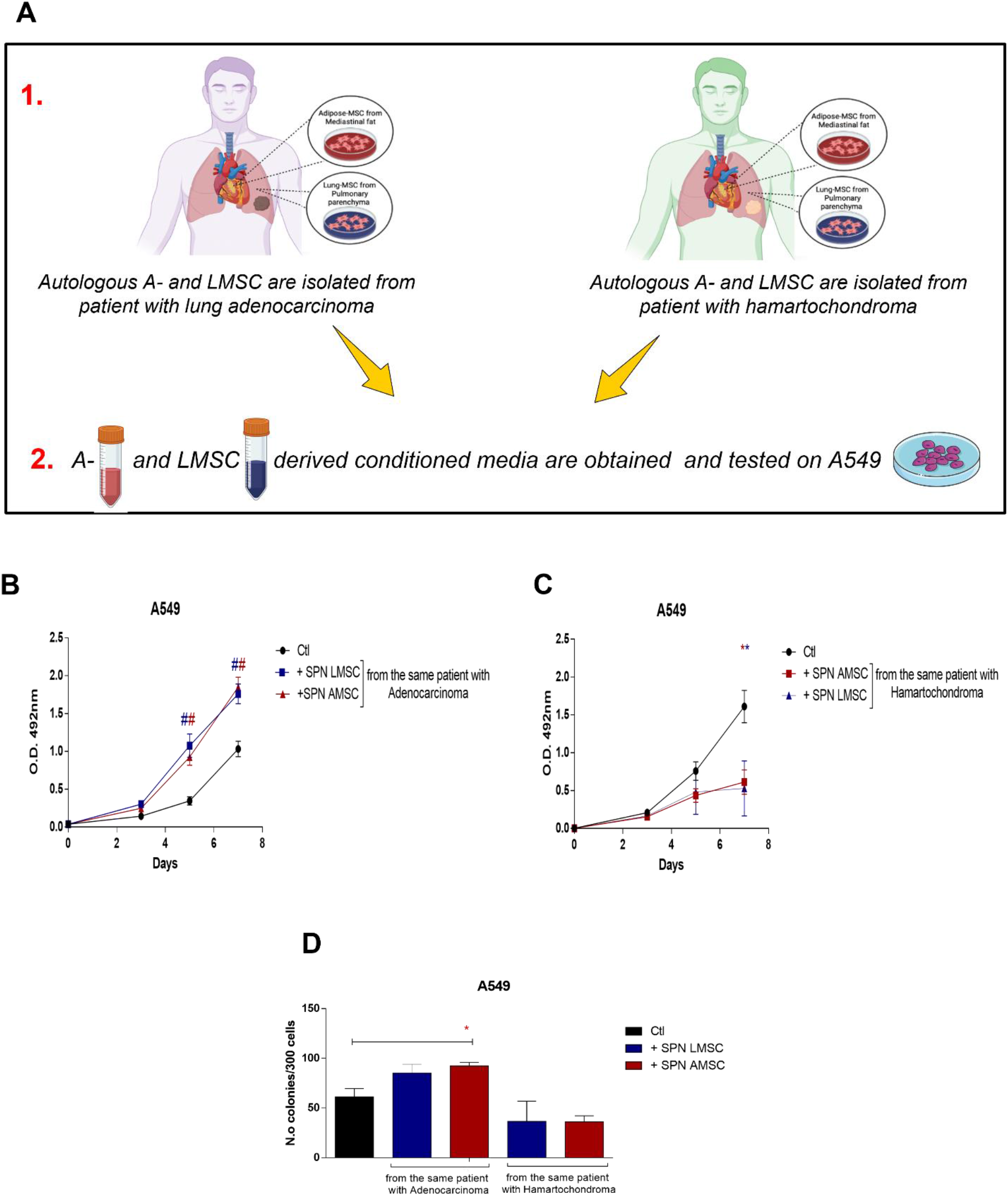
**(A)** Experimental design of the study on A549 cell line. 1 and 2 represent the steps of the experiments. **(B)** A549 proliferation by MTS assay by employing supernatants of autologous A- and LMSC derived from patient with lung adenocarcinoma or **(C)** hamartochondroma. **(D)** Clonogenic assay on A549 with the corresponding autologous supernatants of B and C. N=4 different conditioned media for each source of A- and LMSC. Samples were normalized on time 0. *p<0,05; #p<0,001.

Afterwards, we verified whether the same effects were also reproducible on early-stage human primary lung adenocarcinoma cells (4 lines of LAC staging T1/N0/M0 G1), by employing the same supernatants as for the A549 cell line. The experimental sequence is described in Figure 2A. Notably, both supernatants from autologous LMSC and AMSC (because derived from the same subject) of patients with lung adenocarcinoma were able to sustain cell proliferation of LAC cells similarly to controls at all time points (Figure 2B, p>0.05). Differently, conditioned media of autologous L- or AMSC obtained from patients with pulmonary hamartochondroma, were able to decrease cell proliferation of LAC compared to controls at day 5 and 7 (Figure 2C, day 5 LMSC p<0.001, AMSC p<0.05 *vs* controls; day 7 both LMSC and AMDC p<0.001 *vs* controls). Coherently, we also found a significant enhancement of the clonogenic ability of LAC after the culturing with supernatants of autologous LMSC o AMSC derived from patients with early-stage lung adenocarcinoma (Figure 2D, p=0.04 LMSC and p=0.02 AMSC *vs* controls). Conditioned media derived from autologous L- or AMSC of patient with hamartochondroma decreased or did not alter the clonogenic capacity of LAC cells (Figure 2D, p=0.011 LMSC *vs* controls).

**Figure 2A-D.**
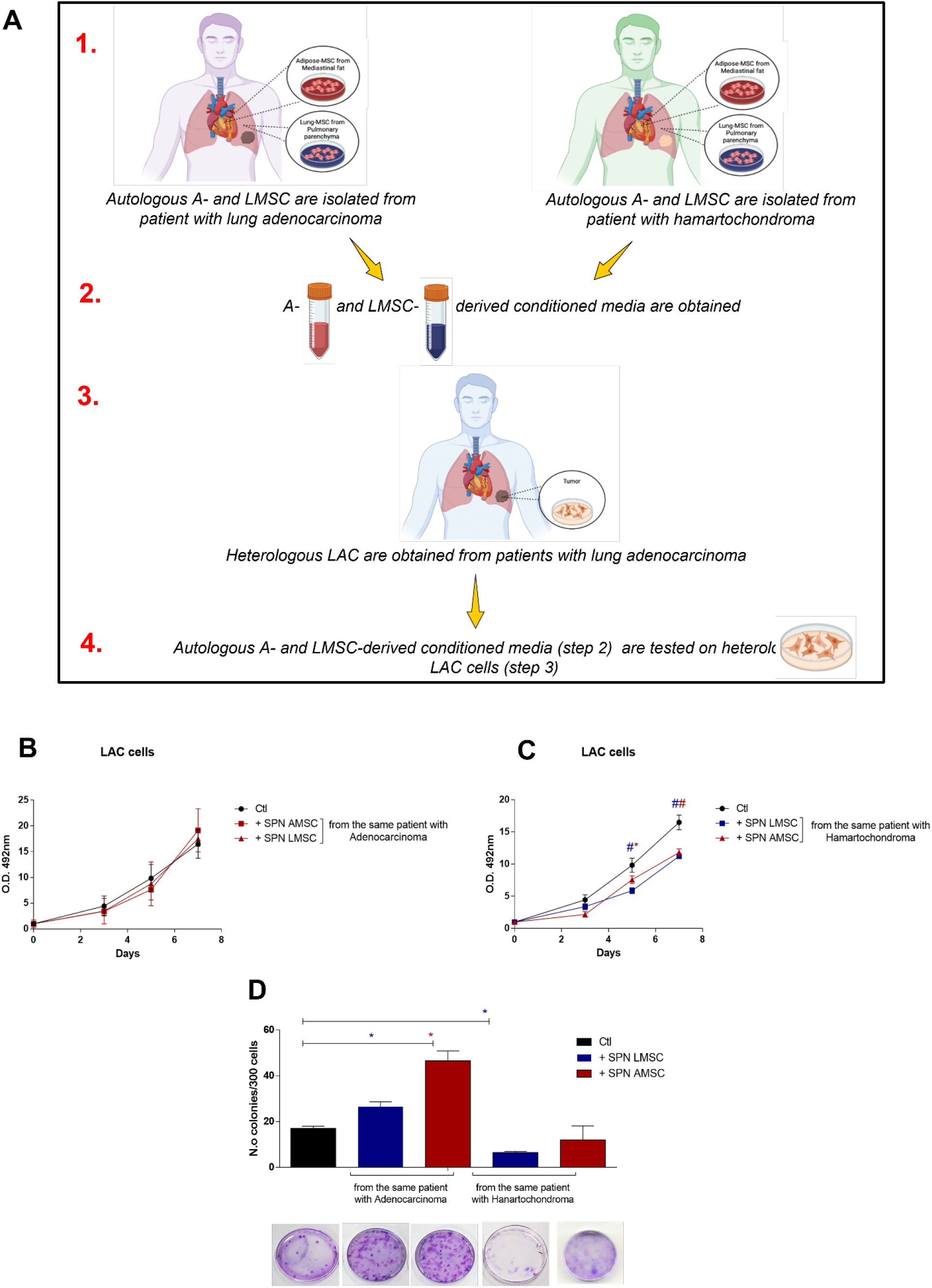
**(A)** Experimental design of the study on human primary lung adenocarcinoma cells (LAC). From 1 to 4 are represented the steps of the experiments **(B)** LAC proliferation by MTS assay with supernatants of autologous A- and LMSC derived from patient with lung adenocarcinoma or **(C)** hamartochondroma. **(D)** Clonogenic assay of heterologous LAC with the corresponding autologous supernatants of B and C. Below the graph representative images of Giemsa staining of the clones generated by heterologous LAC cells after 3 weeks of culture in presence of autologous A- or LMSC-derived conditioned media. N=4 different conditioned media for each source of A- and LMSC. *p<0,05; #p<0,001.

We then evaluated if these effects entailed also a different paracrine signature of autologous MSC-derived microenvironments. Thus, we screened a panel of 80 pro- and anti-inflammatory cytokines and growth factors excreted by both sources of MSC in the conditioned media. Results analysed through the heatmap (soluble factors with no expression in both L- and AMSC were ruled out) revealed that the secretome of both autologous L- and AMSC of patients with early-stage lung adenocarcinoma was similar (Figure 3A), therefore suggesting a comparable in vitro microenvironment. Interestingly, supernatants derived from AMSC of patients with hamartochondroma exhibited a more heterogenous profile of soluble factors respect to that derived from the corresponding autologous MSC counterpart in the lung (Figure 3B). Afterwards, we further examined these secretomes. By keeping constant the source of the stromal fraction during the analysis, we calculated the increasing ratio of the soluble factors between malignant and benign microenvironments. An upregulation ratio of >1.2 was set as cut-off. Then, we sought for those common cytokine or growth factors upregulated among the two microenvironments and we generated a functional protein association network by the STRING software. Results showed a shared upregulation of 6 soluble factors including EGF, IL-1β, IL-3, TNF-α, CCL2, and SPP1 (osteopontin). Notably, the STRING analysis which identifies protein-protein interaction, also displayed that all cytokines but SPP1 were strictly interconnected in the same cluster with a high confident score of 0.7. Afterwards, we used Metascape for the pathway enrichment analysis of this network, and we found that the most significant terms within the cluster consisting of EGF, IL-1β, IL-3, TNF-α, CCL2 were associated to lung fibrosis and pro-inflammatory/fibrotic mediators (Figure 3D).

**Figure 3A-D.**
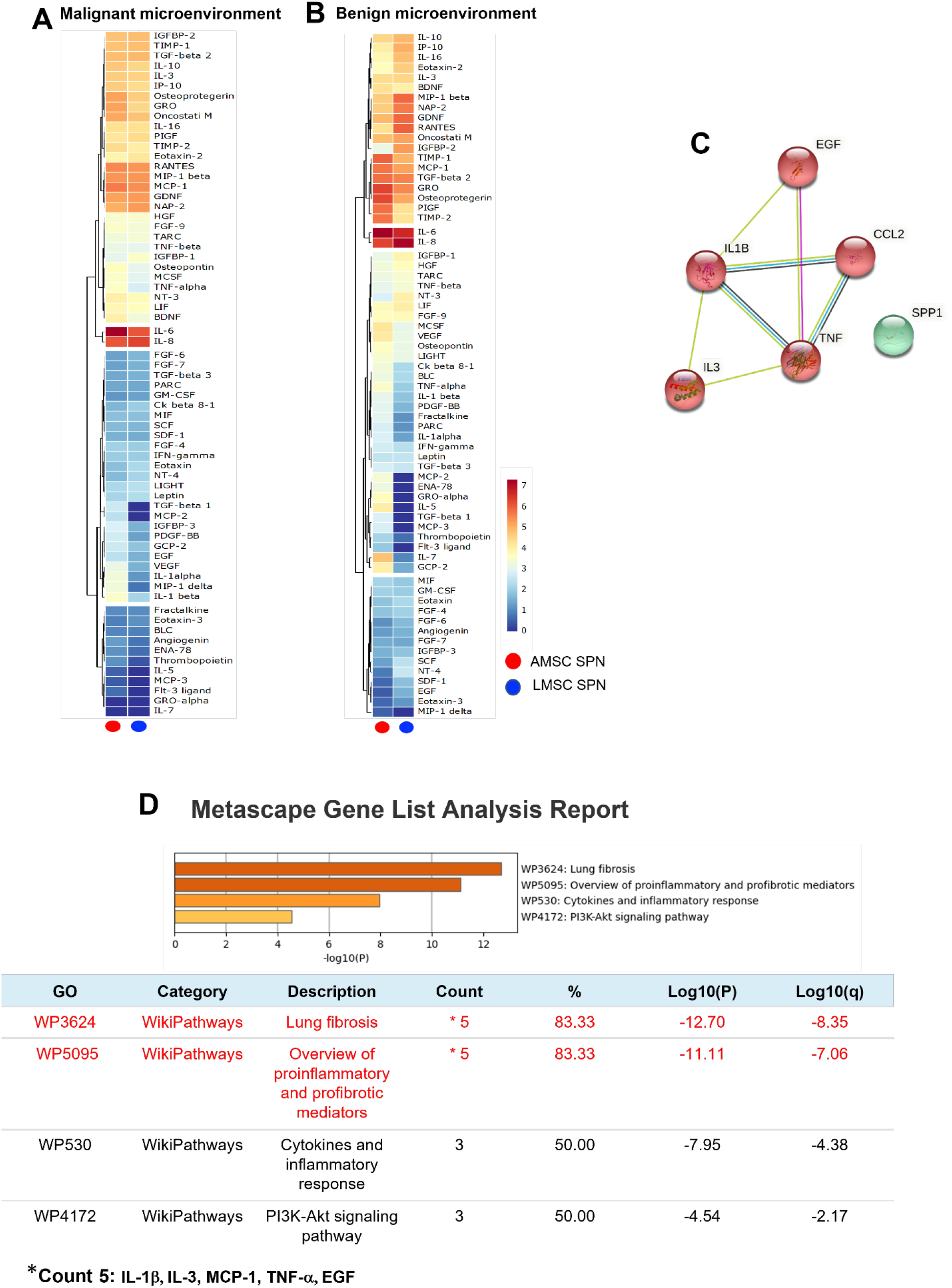
**(A)** Hierarchic clustering heatmap based on the cytokine arrays of autologous A- and LMSC derived conditioned media from patients with lung adenocarcinoma and hamartochondroma (malignant and benign microenvironments, respectively). The red-yellow and the blue range colours indicate cytokines with high and low average levels, respectively. **(C)** Protein-protein interaction network generated by STRING database on the 6 upregulated cytokines commonly shared by autologous A- and LMSC when their malignant and benign microenvironments are compared. The number of nodes is proportional to a strict correlation among the proteins analysed. **(D)** The analysis of the pathway and process enrichment evaluation derived from the Metascape analysis of the cytokines displayed in **(C)** showing that IL-1β, IL-3, MCP-1, TNF-α, EGF are all related to lung fibrosis and inflammation.

These results indicated a potential biological supporting tumor behaviour of AMSC localized in remote areas even at early stages of the disease. Thus, to validate this aspect, we attempt to similarly “educate” in vitro the benign AMSC obtained from patient with hamartochondroma towards a biological “malignant-like” behaviour, by reversing the experiment and culturing benign AMSC for 7, 14 and 21 days with the sole supernatants produced by LAC cells. Afterwards, the same conditioned media produced by AMSC after conditioning with adenocarcinoma, was removed and replaced on A549 cells which were tested for cell proliferation. We found a significant increase in cell proliferation of A549 at 14 and 21 days of tumour pre-conditioning compared to day 7 (Figure 4, p<0.001 both).

**Figure 4.**
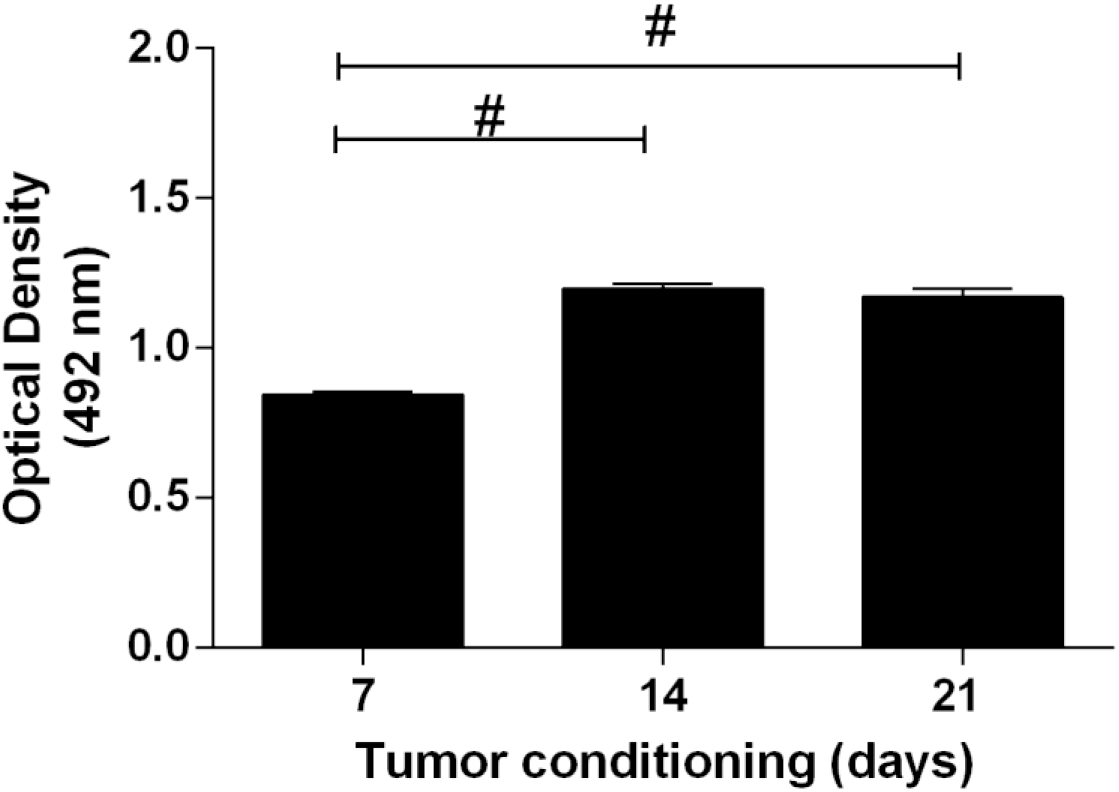
Proliferation assay of A549 after culture with supernatants of benign AMSC previously preconditioned with LAC-derived conditioned media up to 21 days and so retested on A549. The graph shows an increase in cell proliferation of A549 at 14 and 21 days compared to 7 days. Samples were normalized on basal media of AMSC. #p<0,001.

Considering the analysis of the malignant and benign secretome of A- and LMSC (Figure 3A-D) and that miRNAs have been demonstrated as eligible candidate to mediate a long-distance paracrine effect, to instruct MSC and endothelial cells in lung cancer [33], we performed a computational analysis using the miRNA-target prediction tools miRecords, miRTarBase and TarBase combined to a revision of the literature on lung cancer of miRNAs targeting the pool of the 5 cytokines. We found four miRNAs including miR-126, 101, 486 and let-7-g. To validate the four miRNAs, we explore their expression levels on matched lung cancer tissues and samples blood from the same set of patients employed to isolate LAC cells. Results showed that among all analysed miRNAs the miR-126 displayed the highest expression in the lung cancer tissue (Figure 5A, p<0.01 and p<0.001), differently from the circulation where is the miR-486 to show the highest levels (Figure 5B, p<0.05 and p<0.01).

**Figure 5A-B.**
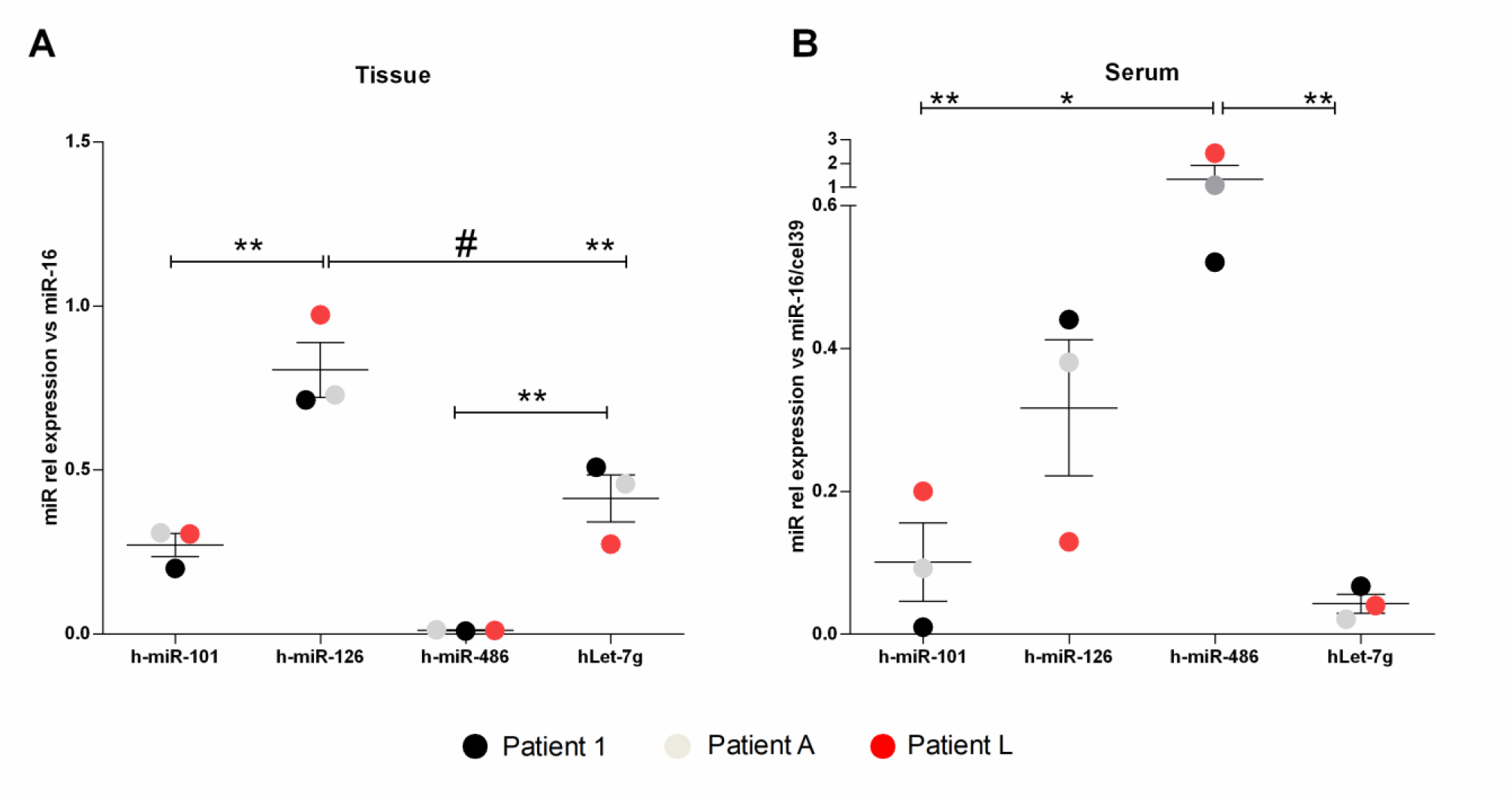
Real Time PCR for miR-101, 126, 486 and let-7-g in **(A)** cancer tissues and the corresponding **(B)** sera of 3 patients with lung adenocarcinoma at early stages. Samples were normalized on miR-16 and miR-16/cell39 for tissue and sera, respectively. **p<0,01; #p<0,001.

## 4. Discussion

This short report highlights how the tumor microenvironment is already defined at early stages, such that the stromal fraction can be influenced even at remote sites and in absence of metastasis. Intriguingly, remote AMSC derived from subjects with lung adenocarcinoma, are permissive to cell proliferation and clonogenic properties when tested on both A549 and LAC cells similarly to the stromal lung counterpart. Oppositely, this effect is lost in “benign conditions”. The first important point of novelty of our brief study is that we have been able to isolate both A- and LMSC within the same patient (autologous stromal cells), therefore ruling out the variability in biological performance within the individual. A further point of originality is that our study has been focused on the early stages of lung adenocarcinoma, which have been currently given more clinical attention. Accordingly, we have shown that at early stages of the tumor, malignant-like microenvironments are already generated from AMSC at remote sites, and they can be overlapped to the lung stromal counterpart, which is tumor adjacent. This is in line with the concept that cancer cannot be interpreted only as a local disease, but rather than a systemic disorder [34,35] and with the concept of tumor permissive microenvironment as the result of the interaction between stroma and cancer [36]. Our data extend this idea also to early stages of tumor and not only when metastasis occurs [37]. In fact, several evidence already exists regarding the cancer cell spreading, considered as a very early event [38].

The biological alterations we have highlighted here, are centred on the microenvironment, which is considered a critical hallmark to elucidate mechanisms of cancer plasticity [39] and where the stromal fraction exerts a critical regulatory role within the tissue [40]. Specifically, our data show that difference among the secretome obtained by benign and malignant-like microenvironments of AMSC at early stages, is limited to the increase of a small pool of soluble factors. Notably, among them (IL1-β, IL-3, MCP-1, TNF-α), the EGF, the most acknowledged target for lung cancer therapy [41], emerges. We also found that this set of soluble mediators is functionally interconnected and related to lung inflammation and fibrosis. The grade of fibrosis in lung cancer is a key issue and represents the modification of a permissive microenvironment induced by the continuous crosstalk between tumor and cancer-associated fibroblasts (as part of the stromal fraction), which leads to the manipulation of the extracellular matrix components and to the transition towards the epithelial-mesenchymal traits [42]. Lung fibrosis may be a prerequisite for the development of lung adenocarcinoma [43] and so is the perpetuating inflammatory condition which fosters the most suitable biological background for tumor progression [44]. Pro-fibrotic markers (alpha-smooth muscle actin, fibrillar collagens, SMAD3) expressed in histological samples of patients with lung cancer, are correlated to low survival [45]. Other important observations derived from pulmonary idiopathic fibrosis cases and the interstitial fibrosis, considered as an independent risk variable for lung adenocarcinoma [46-48]. Lung fibrosis also positively correlates with a glycolytic metabolism of the tumor in subjects with IIIA NSCLC [49].

Notably, we have provided a first biological indication of the possibility to “educate” the benign AMSC towards a malignant-like behaviour. This is in line with several observations regarding the process of educating MSC [50] which has been described for bone marrow derived MSC differentiating towards malignant phenotype, once recruited by cancer microenvironment [51]. Novel clinical applications by employing chemotherapic agents or enhancing CAR-T cells/-NK in cancer immunotherapy, exploits the ability to educate MSC to guide the tropism of the stromal fraction [52,4]. Our study is also coherent with additional reports showing that tumor cells educate MSC depending on tumor microenvironment [53], strengthening the significance of the microenvironmental control exerted by cancer.

Although we have not identified the exact molecular mechanism by which a malignant microenvironment can be also generated in remote stromal area, we have provided a first biological correlation between the secretome (the pool of the 5 increasing soluble factors between malignant and benign microenvironments) of the stromal fraction at remote sites and changes of matched circulating and tissue miRNAs. Our data highlight how serum levels of miR-101, 126 and let-7g already reflect the same expression profile of the corresponding cancer tissue in patients with early stages of lung adenocarcinoma. The miR-486 represents an exception in our findings as its levels are downregulated and upregulated in tissue and serum, respectively. This is not surprising, considering that the decrease of miR-486 found in NSLC tissues, is inversely correlated to both lung metastasis [54] and cancer stages [55]. Plasma levels or miR-486 are reported to increase after NSLC resection [56]. Based on these findings, miR-486 is currently considered as one of the most significant prognostic marker for early diagnosis in NSLC [57]. Besides, the miR-126 which is known as angiomiRNA and mainly produced by platelets and endothelial cells [31], is already upregulated in both cancer tissue and serum, likely suggesting a potential early involvement of a dysregulated angiogenesis towards the influence of lung adenocarcinoma at early stages.

## 5. Conclusions

Our study has several limitations. We have not demonstrated the direct association between the soluble factors-miRNAs and systemic effect towards stromal cells at remote sites upon the influence of lung adenocarcinoma at early stages. The education of the stromal fraction cannot be also restricted to the sole secretome by cancer cells and certainly additional molecular and biological mechanisms including genomic alterations and mutations need to occur, in order to favour the progression of lung cancer. Besides, we must consider that AMSC retain the intrinsic ability to favour cell proliferation and proangiogenic properties depending on the adipose depot [20], suggesting further prudence to the use of MSC in cancer-related clinical applications.

Despite this, our study sheds a further light on the complex relationship between cancer and stromal compartment mediated by the microenvironment to communicate with niches at remote sites.

## Data Availability

Main data generated or analysed during this study are included in this article, and detailed data are available from the corresponding authors on reasonable request.

## Ethical Approval

The methodology described in this study has been conducted in compliance with the tenets of the Declaration of Helsinki for experiments involving human tissues. This study was approved by the Ethical Committee of S. Andrea Hospital, Rome (Prot. CE N. 5195/2013). Medical records of each patient were collected in a clinical database. All data were de-identified and analysed anonymously.

## Authors’ contributions

EDF wrote the manuscript and conceived the study, AB cell cultures, XD performed informatic analysis, PR performed statistics, AV histology, CM, ER and MI surgery, AC conceived the study and revised the manuscript.

## Acknowledgments

We thank the Department of Medical-Surgical Science and Biotechnologies based in Latina for the continuous support

